# Insularity forcing on plant persistence strategies in edaphic island systems

**DOI:** 10.1101/2021.05.13.444066

**Authors:** Luisa Conti, Francisco E. Méndez-Castro, Milan Chytrý, Lars Götzenberger, Michal Hájek, Michal Horsák, Borja Jiménez-Alfaro, Jitka Klimešová, David Zelený, Gianluigi Ottaviani

**Author notes:** **Corresponding author:** Luisa Conti, Faculty of Environmental Sciences, Czech University of Life Sciences, Prague. **Biosketch** LC is a plant ecologist, mainly interested in abiotic, biotic, and geographical drivers of plant assembly. She has also explored drivers of species temporal stability and ecosystem functions, as well as invasions dynamics and their impacts. Particularly, she uses plant functional patterns to unravel these ecological processes. More recently, she has also been investigating remote sensing approaches for quantifying taxonomical and functional diversity. https://www.researchgate.net/profile/Luisa-Conti.

## Abstract

**Aim:** Trait-based approaches are increasingly implemented in island biogeography, providing key insights into the eco-evolutionary dynamics of insular systems. However, what determines persistence of plant species once they have arrived and established in an island remains largely unexplored. Here, we examined links between non-acquisitive persistence strategies and insularity across three terrestrial edaphic island systems, hypothesising that insularity promotes strategies for local persistence.

**Location:** Europe: Western Carpathians, Moravia, and Cantabrian Range.

**Time period:** Present.

**Major taxa studied:** Vascular plants.

**Methods:** For each system, we used linear models at the island scale to test whether persistence-related plant trait patterns (average trait values and diversity) depend on three insularity metrics (island size, isolation and target effect). We focused on patterns of edaphic island specialists because, in contrast to matrix-derived species, their presence is confined to the edaphic islands.

**Results:** We found that insularity metrics explained large proportions in the variation of the average and diversity of persistence-related traits of edaphic island specialists. Insularity was associated with a decline in the proportion of island specialists that have clonal abilities, yet it affected trait values of specialists towards enhanced abilities to persist locally (e.g. more extensive lateral spread) while reducing trait variability. Higher degrees of insularity within the systems were translated to stronger effects on functional trait patterns.

**Main conclusions:** Insularity affects plant species diversity, distribution and forms in terrestrial island-like systems, similarly as it is assumed for true islands. Insularity – measured using a single (island size, isolation) or combined (target effect) predictors – may operate selecting for enhanced and less diverse persistence strategies. Ultimately, this process, which we call insularity forcing, operates as a selective process to promote species ability to avoid local extinction and to persist on terrestrial islands.

## Introduction

Plants have evolved a myriad of ecological strategies to survive in a vast range of environmental and geographical configurations (Díaz et al., 2016). One vital set of strategies are those related to the ability to persist in a given area, which allows plants to maintain viable populations and avoid local extinction (Klimešová, Martínková, & Ottaviani, 2018). These strategies can be described by different plant traits linked to specific functions, such as investment into sexual and/or vegetative reproduction, dispersal of propagules, resprouting after disturbance, and space occupancy (and captured by clonal, bud-bank, seed traits and life forms, Klimešová et al., 2018; Ottaviani et al., 2020; Saatkamp et al., 2019; Violle, Navas, Vile, Kazakou, & Fortunel, 2007; Weiher et al., 1999) While some of these functions are better studied, e.g. those associated with dispersal (Levin, Muller-Landau, Nathan, & Chave, 2003), others, mainly those linked to clonal and bud-bearing organs, are regularly overlooked (Klimešová et al., 2018, 2019; Ottaviani, Martínková, Herben, Pausas, & Klimešová, 2017). This is because sampling of these traits is generally more laborious and time-consuming (being associated with the belowground location of most organs) compared to more commonly used aboveground traits (e.g. leaf economics; Wright et al., 2004). This gap is common in plant ecology research at large, but it is particularly striking in studies focused on insular systems, where long-distance dispersal events are rare, and thus persistence strategies are key for ensuring population maintenance and avoid extinction (Ottaviani et al., 2020).

Compared to non-insular environments, insular systems often host endemic-rich, genetically and eco-morphologically distinct biota (Darwin, 1859; MacArthur & Wilson, 1963, 1967; Burns, 2019; Carlquist, 1974) generated by the eco-evolutionary history associated with a limited spatial extent (König et al., 2021; Taylor, Weigelt, König, Zotz, & Kreft, 2019). Insular systems are not restricted to oceanic islands but also include terrestrial island-like systems such as inselbergs, edaphic islands, and mountaintops (Harrison, Safford, Grace, Viers, & Davies, 2006; Henneron, Sarthou, de Massary, & Ponge, 2019; Jiménez-Alfaro, García-Calvo, García, & Acebes, 2016). For plant specialist species of terrestrial island-like systems, these distinct patches in the landscape may operate as islands because the surrounding matrix is generally inhospitable to their establishment, similarly to water for true islands (Horsák et al., 2012; Ottaviani et al., 2020). Because of the specific spatial configuration of insular systems, island biogeographic studies have typically focused on colonisation (and related dispersal abilities; Franklin et al., 2013; Taylor et al., 2019) and speciation processes (MacArthur & Wilson, 1967; Warren et al., 2015). Less is known about persistence, which can be considered as the opposite of local extinction or extirpation (Auffret, Aggemyr, Plue, & Cousins, 2017). Moreover, trait-based approaches that focus on persistence abilities are virtually non-existent, and as a result, plant persistence strategies and related processes remain poorly understood in insular systems (Ottaviani et al., 2020).

Insularity effects depend on two main components, namely island size and isolation (MacArthur and Wilson, 1963; Lomolino, 2000; Mendez-Castro et al., in press). Island size is considered a key determinant of species richness, with larger islands typically containing more species than smaller ones (Lomolino, 2000b; Lomolino & Weiser, 2001). Also, larger islands tend to have higher variation in geology, topography, and microclimate (hence habitat diversity), which can also contribute to higher species richness (Hortal, Triantis, Meiri, Thébault, & Sfenthourakis, 2009; Keppel, Gillespie, Ormerod, & Fricker, 2016). Spatial isolation relates mainly to species colonisation chances: islands located more distantly from their species source (i.e. mainland or neighbouring islands) are likely to receive fewer species than islands located closer to the species source (Carter, Perry, & Russell, 2020; Warren et al., 2015). At the same time, species on more isolated islands, both true islands and terrestrial islands, are exposed to a higher risk of local extinction (Jiménez-Alfaro et al., 2021; Warren et al., 2015; Whittaker, Fernández-Palacios, Matthews, Borregaard, & Triantis, 2017). Most research in recent years has focused on disentangling and quantifying the relative contribution of island size and isolation in shaping species diversity patterns (e.g. Weigelt and Kreft, 2013; Jacquet et al., 2017). This effort has provided many valuable insights for island biogeography, yet sometimes the effect of insularity might be only detected by considering the joint effect of island size and isolation working together (e.g. by the target effect) (Gilpin and Diamond, 1976, Mendez-Castro et al., in press).

The effect of insularity on island plants should be reflected in distinct patterns of functional traits (Biddick, Hendriks, & Burns, 2019; Burns, 2019; Carlquist, 1974; Taylor et al., 2019). By examining trait patterns, we can gain insights into different plant strategies and processes occurring in insular systems, including those promoting or hindering persistence (Auffret et al., 2017). In this work, our primary aim is to explore whether and how strongly plant persistence strategies (captured by clonal, bud-bank, seed traits and life forms) are linked to insularity – to our knowledge, the first attempt of this kind. We do so by examining three steadily isolated terrestrial island-like systems in Europe characterised by distinct edaphic conditions contrasting with the surrounding landscape (hereafter edaphic islands). We focus particularly on the persistence strategies of plant specialist species of these systems.

Based on a recent overview of the topic (Ottaviani et al., 2020), we expect that enhanced abilities of plants to persist locally will be related to a higher degree of insularity – in terms of the individual and joint effect of island size and spatial isolation. Specifically, we predict that with higher degrees of insularity, plant specialists of edaphic islands are distinguished by: (H1) trait values and functional strategies indicating better ability to persist locally, such as higher percentage of clonal species, longer-lived connections between ramets, higher capacity of lateral spread, larger and better-protected bud-bank traits, and heavier seeds; and (H2) less diversified persistence strategies, i.e. species sharing similar functional trait values (reduced variability).

## Methods

In brief, to test whether insularity drives persistence-related trait patterns, we used vegetation data from three different edaphic island systems and joined them with functional trait data obtained from available databases. For all the three systems, we computed, at the island level, trait averages and functional diversity, as well as insularity metrics, namely island size, isolation, and target effect (the ratio between isolation and island size; Mendez-Castro et al., in press). We used linear models to explore the relationship between single functional indices and each individual insularity predictor.

### Edaphic island systems and species selection

We examined three different types of terrestrial island-like systems experiencing long-term isolation. Namely, these are spring fens in the Western Carpathians, Czech Republic and Slovakia (hereafter fens), rocky outcrops in the Třebíč region in Moravia, Czech Republic (hereafter outcrops), and siliceous mountaintops in the Cantabrian Mountains, Spain (hereafter mountaintops) (Figure 1). All the three systems are distinguished by patches of distinct soils supporting specific vegetation types, which are different from the surrounding landscapes. Therefore, they are considered edaphic islands. The target vegetation types were i) treeless calcareous fens; ii) dry, shallow-soil acidophilous grasslands for outcrops; and iii) alpine grasslands on acidic bedrock for mountaintops. The three systems, which differ in their geographical extent and environmental conditions, comprise floras with different species composition and species richness. An expert-based categorisation was carried out for identifying specialist species (and matrix-derived species). The total number of specialist species was 57 for fens, 29 for outcrops and 42 for mountaintops, whereas the total number of matrix-derived species was 260 for fens, 175 for outcrops and 125 for mountaintops.

**Figure 1.**
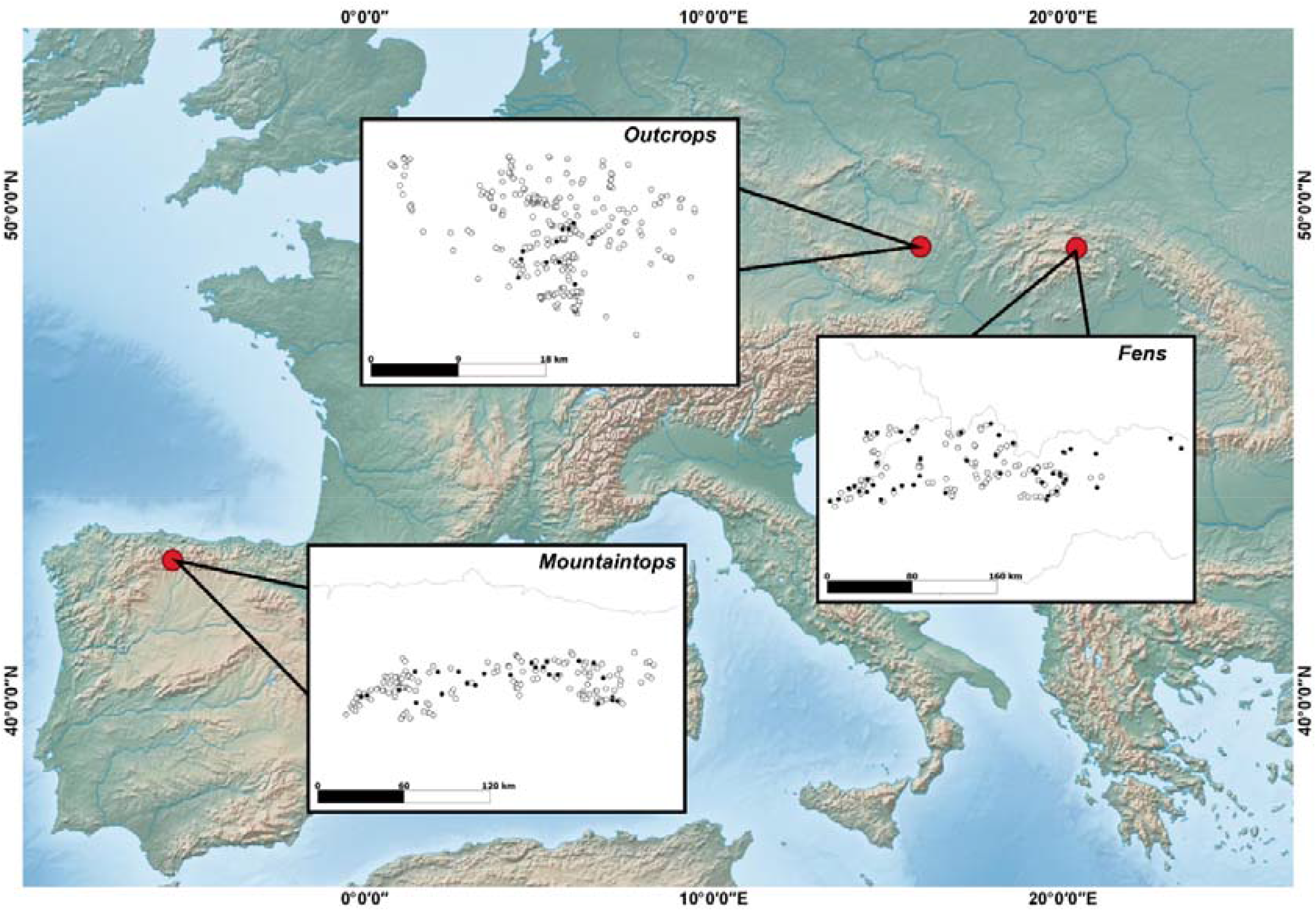
Geographical location and landscape configuration (inset) of the three terrestrial edaphic island systems in Europe. Black circles indicate the sampled habitat islands, whereas empty circles represent the other (not sampled) islands in the surrounding landscape.

For fens, one 4 × 4 m vegetation plot was sampled at the centre of each fen, which generally corresponds to the most homogeneous and best-preserved part of the edaphic island. In addition, a full species list for each island was compiled by a census across each of the edaphic islands so that a full species list for each island was assembled (Hájek et al., 2011; Horsák et al., 2012). For outcrops, vegetation data were collected by performing plot-based surveys (4 plots of 0.5 × 0.5 m at each island) and, also in this case, complemented by a census. For mountaintops, a total of 284 vegetation plots with sizes between 10 and 50 m^2^ were sampled on the target vegetation type (with varying number of sampling plots, proportional to island size). We sampled 49 islands for fens, 20 islands for outcrops, and 25 islands for mountaintops. In all systems, we considered only species occurrence (presence or absence data) at the island level in the analyses.

### Persistence-related trait data and functional metrics

For all the species, we compiled functional trait data relative to persistence strategies from databases, such as Pladias (Chytrý et al., 2021), CLO-PLA3 (Klimešová, Danihelka, Chrtek, de Bello, & Herben, 2017), BiolFlor (Klotz, Kühn, & Durka, 2002) and LEDA (Kleyer et al., 2008). We gathered information on whole-plant strategies related to reproductive type and belowground clonal and bud-bearing organs (Klimešová et al., 2019; Ottaviani et al., 2020; Ottaviani et al., 2017; Weiher et al., 1999; Westoby, 1998) (Table 1). For the mountaintops’ dataset, the number of species for which trait data was available was very low. Therefore, information on the clonal ability and life form of mountaintop species was gathered directly from field observations. Information on the reproductive type (i.e. whether a species can reproduce only sexually, only vegetatively or both) was used to calculate the percentage of clonal species on each edaphic island. We considered as clonal all species that can reproduce vegetatively. Species data summary and trait-data completeness within each study system are presented in Supporting Information Appendix S1.

**Table 1.** List of persistence-related traits included in this study. Trait definition, variable type, units, specific non-acquisitive functions, and source database are reported. Key references related to databases, and trait definitions and their ecological relevance are in the main text.

| Trait name | Definition, variable type and units | Specific function | Source |
| --- | --- | --- | --- |
| Raunkiaer life form | Categories identified based on bud location in the plant body in relation to the soil surface (discrete; nominal) | Multifunctional persistence strategy | Pladias, Field observations |
| Reproductive type | Whether a plant species is capable of reproducing only sexually, only vegetatively or both – used to calculate the percentage of clonal species on each island (discrete; nominal) | Reproductive type | BiolFlor, Field observations |
| Seed mass | Weight of the seed (continuous; mg) | On-spot persistence; Dispersal ability | LEDA |
| Size of the belowground bud bank | Total number of buds stored in belowground bud-bearing organs per individual (continuous; number) | Resprouting ability (vigor) | CLO-PLA3 |
| Depth of the belowground bud bank | Location of the bud bank below the soil surface (continuous; cm) | Resprouting ability (bud protection) | CLO-PLA3 |
| Type of clonal growth organ | Organs dedicated to clonal growth (discrete; nominal) | Space occupancy; Resource sharing | CLO-PLA3 |
| Persistence of the clonal growth organ | Period of connection between offspring individuals and parental individual in clonal plants (continuous; year) | Space occupancy; Resource sharing | CLO-PLA3 |
| Lateral spread | Distance between offspring individuals and parental individual in clonal plants (continuous; mm/year) | Space occupancy | CLO-PLA3 |

Because we expected the effect of insularity to be more pronounced on specialists, which are confined to the edaphic islands (compared to matrix-derived species), we focused on functional metrics for these species (Horsáková, Hájek, Hájková, Dítě, & Horsák, 2018; Ottaviani et al., 2020). Therefore, we calculated unweighted trait averages and functional diversity metrics for specialist species across the three systems. However, to qualitatively assess differences in trait patterns between specialists and matrix-derived species, we also calculated these metrics for the latter. All results related to matrix-derived species are reported in Appendix S2. For each island, we calculated trait average as the mean value for each quantitative (numerical) trait, namely depth of the belowground bud bank, lateral spread, persistence of the clonal growth organ, seed mass, and size of the belowground bud bank (Table 1). We calculated diversity by using the mean pairwise distance of the species’ traits. For quantitative (numerical) traits, we used Euclidean distance, while for qualitative (discrete categorical) traits (Raukiaer life form, type of clonal growth organ), we used Gower distance (Gower, 1971). In addition, we calculated the multi-trait mean pairwise distance, considering all the traits and using the Gower distance.

### Insularity metrics

We explored the effect of insularity, namely of island size, spatial isolation and their joint effect on plant populations. Hence, we used three insularity metrics which proved to be the most important predictors (tested against other connectivity and network metrics) of edaphic island specialist species richness in the same study systems (unpublished data). In detail, we used 1) edaphic island size (hereafter ‘island size’; m^2^), defined as the area of the target vegetation type present on each island (which can correspond to the entire island area or part of it); 2) distance to nearest species source (hereafter ‘isolation’; m), measured as the distance of the target island to the nearest putative species source – one of the islands in the same system scoring above the third quartile of both its size and plant specialist species richness; and 3) target effect (a dimensionless property of islands generated by scaling the distance to the species sources by the island size), calculated as the natural logarithm of the ratio between isolation and the square root of the island size. High values of target effect indicate a lower chance of colonisation, hence a strong role of isolation and a high degree of insularity. Size and isolation were log-transformed to normalise these variables prior to the statistical analyses.

### Statistical analyses

Because we were interested in exploring how persistence-related trait patterns depend on each insularity metric and because correlation tests on insularity metrics for each system identified collinearity issues (Supporting Information Appendix S3), we used each insularity metric as a single explanatory variable for each functional index in separate models. For each edaphic island system, we explained the variation in each of the functional indices, trait averages (H1) and functional diversity (H2) at the island scale through ordinary least-square linear models. Therefore, for each functional index, we calculated three different linear models, each containing the functional index as the response variable and a single insularity metric as the explanatory variable (predictor). The same models were fitted for the set of functional metrics related to the matrix-derived species (Appendix S2). For each model, we calculated standardised effect sizes and relative standard errors (to compare the effects across single models), coefficient of determination R^2^ (explained variation, to measure the goodness of fit), and p-value (significance of the coefficient). The linearity and homoscedasticity of the relationships were assessed by visually inspecting the model residuals, and we revealed no issues with those assumptions. Here, we focus mainly on strong (R^2^ ≥ 0.10) and robust findings (based on the standard error of the model estimate, and in most cases, significant at p-value < 0.05). However, trait-data coverage for mountaintops was considerably sparser than for the other two systems. Here, data were available only for the percentage of clonal species and life form diversity, which restricted our ability to investigate trait patterns in that system. All statistical analyses were done using the R statistical software (version 4.0.3).

## Results

We found significant functional patterns depending on insularity metrics across the three systems under study. However, the number and strength of the relationships varied depending on the type of index and the particular system. Generally, all the patterns found for specialist species were stronger and more numerous than matrix-derived species (Supporting Information Appendix S2).

In all three systems, the insularity predictors explained a considerable amount of variability in the models related to the percentage of clonal specialists (with R^2^ up to 0.56; Table 2). Generally, island size and target effect explained a higher proportion of variability compared to isolation. Island size had a positive effect on the percentage of clonal specialists, while isolation and target effect exerted a negative effect.

**Table 2.** Models output for the proportion of clonal specialist species in the edaphic island systems. Each column represents a different model having one single insularity metric as a predictor. Bold text indicates a relevant statistic (significant coefficient and/or R^2^ ≥ 0.10). All predictors are mean-centered and scaled by 1 standard deviation. *** p < 0.001; ** p < 0.01; * p < 0.05.

|  | FENS |  |  | OUTCROPS |  |  | MOUNTAINTOPS |  |  |
| --- | --- | --- | --- | --- | --- | --- | --- | --- | --- |
|  | Island size | Isolation | Target effect | Island size | Isolation | Target effect | Island size | Isolation | Target effect |
| Coefficient | **8.86 \*\*\*** | **-7.66 \*\*\*** | **-9.43 \*\*\*** | **10.38 \*\*** | -5.50 | **-8.89 \*** | **3.95 \*\*** | 2.46 | **-0.04 \*** |
| Standard Error | (1.32) | (1.47) | (1.23) | (3.01) | (3.66) | (3.26) | (1.26) | (1.41) | (0.02) |
| R<sup>2</sup> | **0.49** | **0.37** | **0.56** | **0.40** | **0.11** | **0.29** | **0.30** | **0.12** | **0.16** |

Regarding the other trait average values, we found that lateral spread was explained by all insularity metrics in both fens and outcrops, although with very low coefficients (Table 3). In particular, lateral spread was negatively affected by island size and positively by isolation and target effect. For fens, we also found that the belowground bud bank was deeper in more insular conditions, with island size having a negative effect and target effect having a positive effect on this trait (with a similar pattern also revealed for the size of the belowground bud bank; Table 3). For outcrops, we found that both persistence of the clonal growth organ and seed mass were influenced by insularity, with target effect negatively linked to the persistence of the clonal growth organ and island size negatively affecting seed mass. The amount of variability explained in these models was generally lower compared to the models regarding the percentage of clonal species and those regarding functional diversity (with R^2^ up to 0.26; Table 3).

**Table 3.**
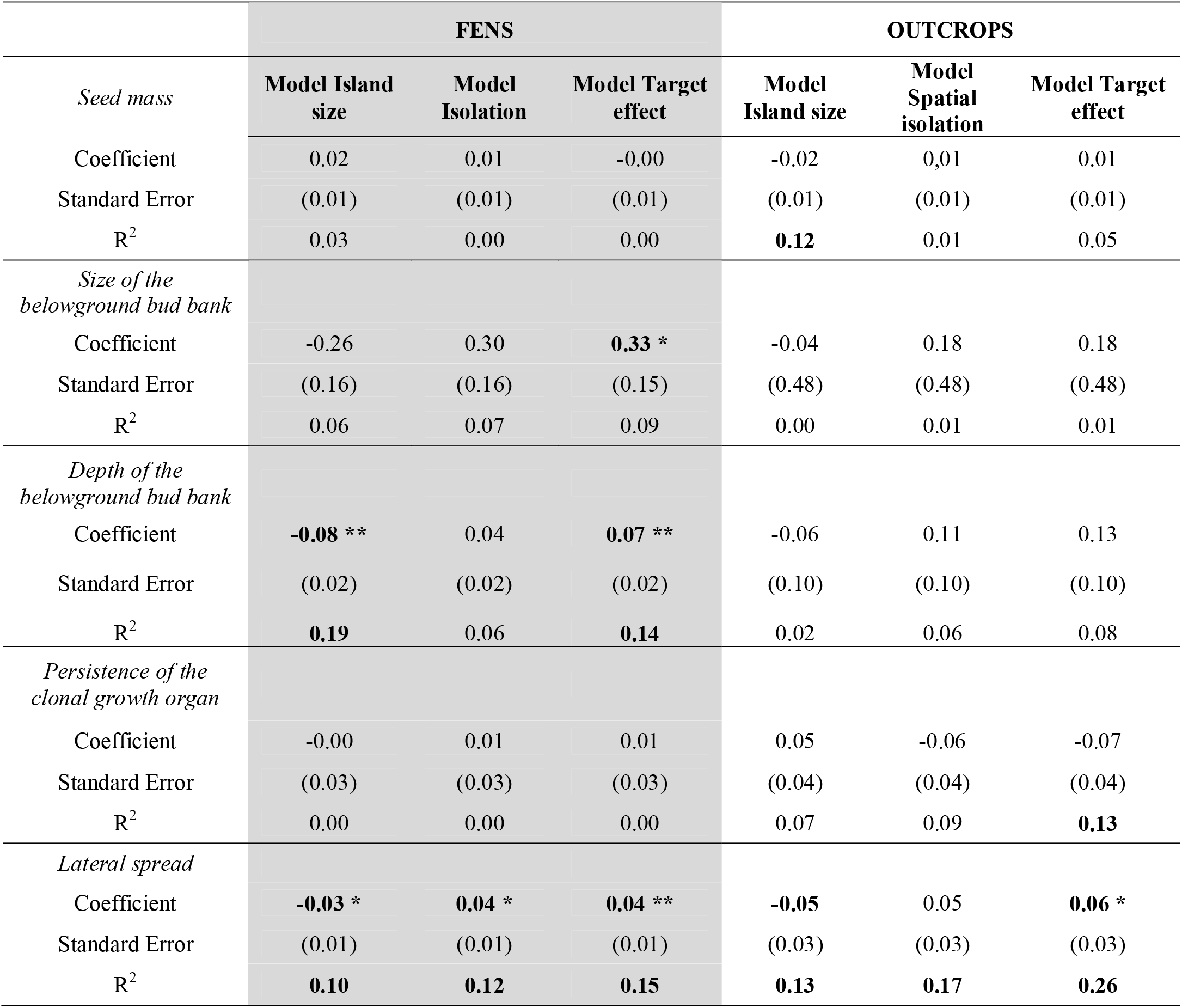
Models output for average functional trait values (trait names in italics). Each column reports a summary for a different model, each having one single insularity metric as a predictor. Bold text indicates an important model output (significant coefficient and/or R^2^ ≥ 0.10). All predictors are mean-centred and scaled by 1 standard deviation. *** p < 0.001; ** p < 0.01; * p < 0.05.

|  | FENS |  |  | OUTCROPS |  |  |
| --- | --- | --- | --- | --- | --- | --- |
|  | Model Island size | Model Isolation | Model Target effect | Model Island size | Model Spatial isolation | Model Target effect |
| *Seed mass* |  |  |  |  |  |  |
| Coefficient | 0.02 | 0.01 | -0.00 | -0.02 | 0.01 | 0.01 |
| Standard Error | (0.01) | (0.01) | (0.01) | (0.01) | (0.01) | (0.01) |
| $R^2$ | 0.03 | 0.00 | 0.00 | **0.12** | 0.01 | 0.05 |
| *Size of the belowground bud bank* |  |  |  |  |  |  |
| Coefficient | -0.26 | 0.30 | **0.33 \*** | -0.04 | 0.18 | 0.18 |
| Standard Error | (0.16) | (0.16) | (0.15) | (0.48) | (0.48) | (0.48) |
| $R^2$ | 0.06 | 0.07 | 0.09 | 0.00 | 0.01 | 0.01 |
| *Depth of the belowground bud bank* |  |  |  |  |  |  |
| Coefficient | **-0.08 \*\*** | 0.04 | **0.07 \*\*** | -0.06 | 0.11 | 0.13 |
| Standard Error | (0.02) | (0.02) | (0.02) | (0.10) | (0.10) | (0.10) |
| $R^2$ | **0.19** | 0.06 | **0.14** | 0.02 | 0.06 | 0.08 |
| *Persistence of the clonal growth organ* |  |  |  |  |  |  |
| Coefficient | -0.00 | 0.01 | 0.01 | 0.05 | -0.06 | -0.07 |
| Standard Error | (0.03) | (0.03) | (0.03) | (0.04) | (0.04) | (0.04) |
| $R^2$ | 0.00 | 0.00 | 0.00 | 0.07 | 0.09 | **0.13** |
| *Lateral spread* |  |  |  |  |  |  |
| Coefficient | **-0.03 \*** | **0.04 \*** | **0.04 \*\*** | **-0.05** | 0.05 | **0.06 \*** |
| Standard Error | (0.01) | (0.01) | (0.01) | (0.03) | (0.03) | (0.03) |
| $R^2$ | **0.10** | **0.12** | **0.15** | **0.13** | **0.17** | **0.26** |

For functional diversity, we found stronger and more numerous significant patterns in fens than in outcrops. We found that for both edaphic island systems, Raunkiaer life forms (also confirmed in mountaintops), type of clonal growth organs, and multi-trait diversity were explained by the insularity metrics (Table 4). Generally, these relationships were positively explained by island size and negatively by isolation and target effect. In fens, diversity of size and depth of the belowground bud bank, and for outcrops, lateral spread were largely explained by all the insularity metrics (with R^2^ reaching up to 0.45, and in most cases between 0.15 and 0.30; Table 4), with a consistent pattern as of those highlighted above (i.e. positive effect of island size, negative of isolation and target effect).

**Table 4.** Models output for functional diversity (trait names in italics). Each column reports a summary for a different model summary, each having one single insularity metric as a predictor. Bold text indicates an important model output (significant coefficient and/or R^2^ ≥ 0.10). All predictors are mean-centered and scaled by 1 standard deviation. *** p < 0.001; ** p < 0.01; * p < 0.05.

|  | FENS |  |  | OUTCROPS |  |  | MOUNTAINTOPS |  |  |
| --- | --- | --- | --- | --- | --- | --- | --- | --- | --- |
| *Raunkiaer*<br>*life form* | Model<br>Island<br>size | Model<br>Isolation | Model<br>Target<br>effect | Model<br>Island<br>size | Model<br>Isolation | Model<br>Target<br>effect | Model<br>Island<br>size | Model<br>Isolation | Model<br>Target<br>effect |
| Coefficient | **0.01 \*\*** | **-0.01 \*** | **-0.01 \*\*** | 0.01 | -0.01 | -0.01 | **0.07 \*\*** | -0.00 | **-0.04 \*** |
| Standard<br>Error | (0.00) | (0.00) | (0.0042) | (0.01) | (0.01) | (0.01) | (0.02) | (0.02) | (0.02) |
| $R^2$ | **0.16** | 0.09 | **0.16** | **0.14** | 0.06 | **0.13** | **0.35** | 0.00 | **0.16** |
| *Seed mass* |  |  |  |  |  |  |  |  |  |
| Coefficient | **0.11 \*\*\*** | **-0.09 \*\*** | **-0.12 \*\*\*** | -0.02 | 0.01 | 0.02 |  |  |  |
| Standard<br>Error | (0.03) | (0.03) | (0.03) | (0.01) | (0.01) | (0.01) |  |  |  |
| $R^2$ | **0.29** | **0.19** | **0.30** | **0.13** | 0.03 | 0.08 | | | |
| *Size of the*<br>*belowground*<br>*bud bank* |  |  |  |  |  |  |  |  |  |
| Coefficient | **0.54 \*** | **-0.68 \*\*** | **-0.73 \*\*\*** | -0.14 | -0.09 | -0.03 |  |  |  |
| Standard<br>Error | (0.21) | (0.21) | (0.21) | (0.43) | (0.43) | (0.43) |  |  |  |
| $R^2$ | **0.12** | **0.19** | **0.22** | 0.01 | 0.00 | 0.00 | | | |
| *Depth of the*<br>*belowground*<br>*bud bank* |  |  |  |  |  |  |  |  |  |
| Coefficient | **0.16 \*\*** | **-0.17 \*\*\*** | **-0.19 \*\*\*** | -0.09 | 0.05 | 0.08 |  |  |  |
| Standard<br>Error | (0.05) | (0.05) | (0.05) | (0.10) | (0.10) | (0.10) |  |  |  |
| $R^2$ | **0.18** | **0.21** | **0.27** | 0.04 | 0.01 | 0.03 | | | |
| *Type of*<br>*clonal*<br>*growth*<br>*organ* |  |  |  |  |  |  |  |  |  |
| Coefficient | **0.02 \*\*\*** | **-0.02 \*\*** | **-0.02 \*\*\*** | **0.01 \*** | -0.01 | **-0.01 \*** |  |  |  |
| Standard<br>Error | (0.01) | (0.01) | (0.01) | (0.00) | (0.00) | (0.00) |  |  |  |
| $R^2$ | **0.22** | **0.20** | **0.28** | **0.28** | **0.12** | **0.27** | | | |
| *Persistence*<br>*of the clonal*<br>*growth*<br>*organ* |  |  |  |  |  |  |  |  |  |
| Coefficient | -0.00 | -0.02 | -0.01 | 0.05 | 0.00 | -0.01 |  |  |  |
| Standard<br>Error | (0.03) | (0.03) | (0.03) | (0.05) | (0.05) | (0.05) |  |  |  |
| $R^2$ | 0.00 | 0.01 | 0.00 | 0.05 | 0.00 | 0.01 | | | |
| *Lateral*<br>*spread* |  |  |  |  |  |  |  |  |  |
| Coefficient | **-0.03 \*** | 0.02 | 0.03 | **0.10 \*\*** | **-0.07 \*** | **-0.11 \*\*** |  |  |  |
| Standard<br>Error | (0.02) | (0.02) | (0.02) | (0.03) | (0.03) | (0.03) |  |  |  |

| R<sup>2</sup> | 0.09 | 0.03 | 0.07 | **0.42** | **0.22** | **0.45** |
| --- | --- | --- | --- | --- | --- | --- |
| *Multiple trait* |  |  |  |  |  |  |
| Coefficient | **0.01 \*\*\*** | **-0.01 \*\*** | **-0.01 \*\*\*** | 0.01 | -0.00 | -0.01 |
| Standard Error | (0.00) | (0.00) | (0.00) | (0.01) | (0.01) | (0.01) |
| R<sup>2</sup> | **0.21** | **0.16** | **0.24** | **0.16** | 0.01 | 0.06 |

## Discussion

Our results confirmed that insularity – measured as island size, isolation or target effect – largely affects functional trait patterns determining plant persistence strategies in edaphic island systems (Klimešová et al. 2019; Ottaviani et al. 2020). Contrary to our expectation, the percentage of edaphic-island specialist species that can reproduce vegetatively declined with insularity. However, we did confirm that specialists experiencing a higher degree of insularity were equipped with functional strategies (in terms of average trait values) that promote their local persistence, such as enhanced abilities to occupy space, higher resprouting vigour after biomass removal (and better-protected buds) and heavier seeds (Jiménez-Alfaro et al., 2016; Rossetto & Kooyman, 2005). Additionally, these strategies appeared to be selected by insular conditions, i.e. more similar trait values shared across species in a given island, generating lower trait diversity. We also found that among the three studied edaphic island systems, the flora of the fen system was most affected by insularity.

### Clonal specialists are less represented in more insular systems, yet they have enhanced abilities to persist locally

We consistently found that the percentage of clonal species specialised to the edaphic islands declined with insularity across the three systems, so rejecting the first part of our H1. This finding may be caused by a functional trade-off between vegetative reproduction and dispersal abilities (Rossetto and Kooyman, 2005; Herben, Šerá and Klimešová, 2015), influencing species occupancy in terrestrial edaphic islands. As a result, clonal species (which account for a large proportion of specialists in all the studied systems: Table S1.2, Supporting Information) seem to be strongly limited in their dispersal. Therefore, clonal specialists are less likely to reach more distant islands than non-clonal specialist species. Conversely, non-clonal species may not follow this rule and rather than constrained by dispersal, their distribution may be influenced by other biotic and abiotic factors. For example, in fens, non-clonal species trait patterns may be largely determined by either biotic interactions, such as competition between their seedlings and bryophytes (Singh et al., 2019), or higher rates of local extinctions due to temporarily hostile environments (Horsák et al., 2012).

The second part of the first hypothesis – expecting that a high degree of insularity promotes persistence strategies – was supported. Specialist species displayed trait values indicative of better abilities to persist locally both in fen and outcrop systems (no data available for mountaintops). These trait values included, for example, a more extensive lateral spread, resulting in a higher ability to occupy the surrounding space and its resources, and deeper buds belowground, thus better protected resprouting tissues. This finding implies that edaphic island specialists in more insular conditions (smaller island size, more pronounced isolation, or larger target effect) tend to employ strategies aimed at maximising local persistence (Jiménez-Alfaro et al., 2016; Kavanagh & Burns, 2014; Rossetto & Kooyman, 2005; Weber, VanDerWal, Schmidt, McDonald, & Shoo, 2014). This can be attained by i) producing heavier seeds (with limited dispersal), ii) better abilities to explore and remain in the proximal area of the parent plant, and iii) enhanced resprouting capacities after biomass removal (Ottaviani et al., 2020). Therefore, insularity does affect functional trait patterns of edaphic island plant species by promoting persistence strategies aimed at successfully maintaining viable populations and, at the same time, preventing their local extinction (Auffret et al., 2017; Ottaviani et al., 2020).

### High degrees of insularity select for similar persistence strategies

We found support for an insularity effect selecting lower variation in trait values, showing that edaphic island specialists tended to share similar persistence strategies (H2). Indeed, higher degrees of insularity consistently reduced the range of persistence-related trait values and strategies in a given island, i.e. functional diversity, similarly to what is predicted for species richness (Ibanez et al., 2018; MacArthur & Wilson, 1967; Warren et al., 2015). This is further reinforced not only by single trait diversity patterns but also by patterns concerning multi-trait and Raunkiaer life form, traits which can be considered proxies for the multifunctional island persistence syndrome, at least for the non-acquisitive dimension examined here (Ottaviani et al., 2020). These findings were further supported by the evidence that these patterns were revealed across all three edaphic island systems, yet with varying strength depending on the degree of the insularity within each system: in fens, there was a stronger and widespread effect on functional diversity patterns (and more marked in specialists species compared to matrix-derived species), whereas in outcrops and mountaintops, this effect was weaker and less pronounced (although our inference power for mountaintops was limited).

### Insularity as a selective force: the way forward and drawbacks

Taken together, our findings suggest that insularity (measured by isolation, island size and target effect) can select traits related to plant persistence, forcing species towards adaptive strategies appointed to prevent local extinction (Auffret et al., 2017). In fact, our work pinpoints how insularity would act as a selecting force by sorting clonal (and non-clonal) species from the broader species pool (according to classic island biogeography predictions); promoting adaptive persistence strategies in terms of trait means (i.e. enhanced abilities to persist locally and therefore avoid local extinction); and fine-tuning those strategies towards lower diversity values (i.e. similar values shared across species); see Figure 2. Indeed, insularity appears to exert its effect on multiple facets of biological diversity, such as trait averages and functional diversity of persistence strategies in edaphic island systems. We propose to call this process “insularity forcing” on island plant forms and functions – in parallel to what is termed “regional forcing” for plant species diversity and composition on islands (Ibanez et al., 2018).

**Figure 2.**
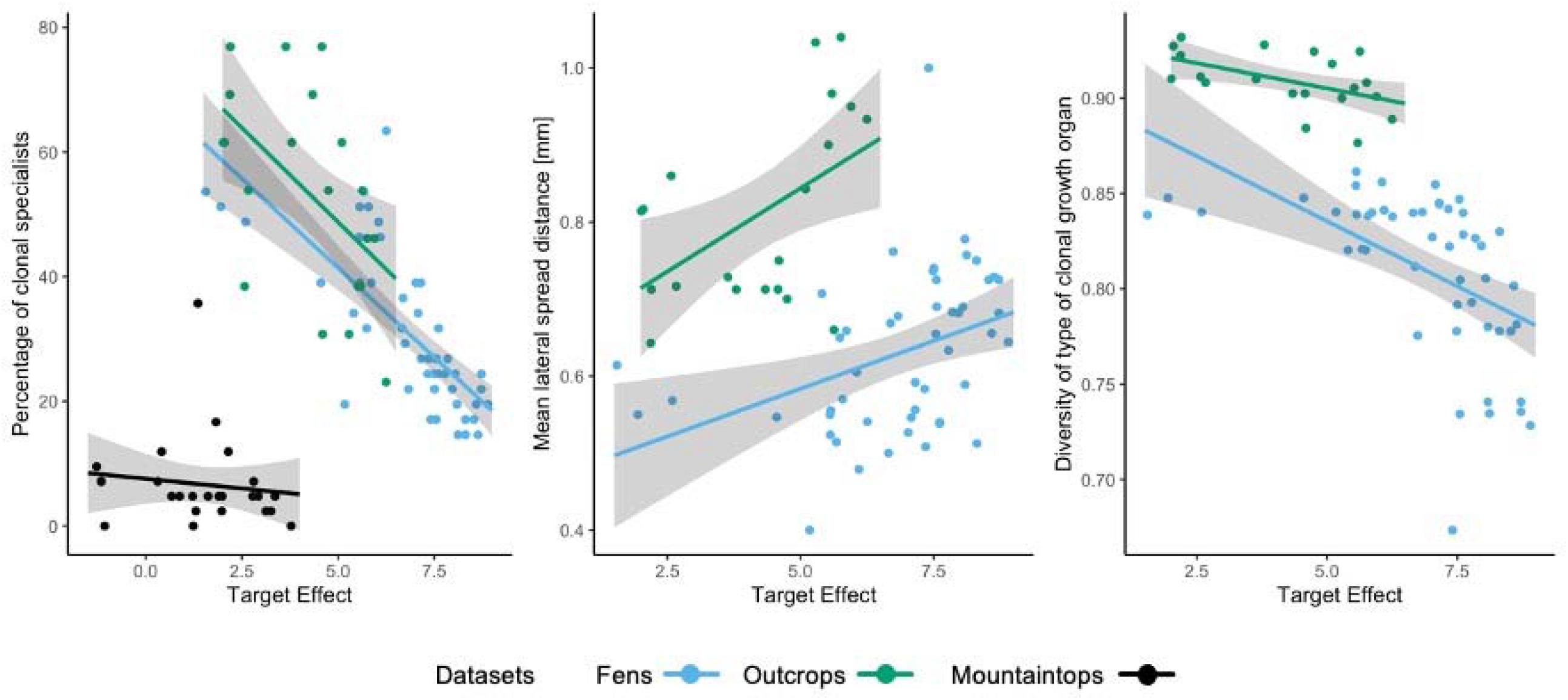
Effect plots of selected models, showing the ‘insularity forcing’, i.e. effect of insularity (target effect) on percentage clonal species, mean trait values (lateral spread distance), and diversity of persistence trait values (type of clonal growth organ) of specialists species in the three edaphic island systems considered.

As any study, our research has its own limitations, and we acknowledge that other abiotic and biotic factors may affect the observed functional trait patterns. In particular, other environmental variables associated with geology, topography and climate may affect habitat diversity, and thereby species composition and plant functional traits (Lavorel & Garnier, 2002), confounding the aforementioned “insularity forcing”. However, having explored the trait-insularity links for both edaphic island specialist and matrix-derived species, we have accounted for possible spurious patterns, and therefore the relationships revealed here can be considered plausible and robust. Yet, future studies may specifically disentangle and quantify the conjoint effect of insularity and other environmental factors on persistence-related functional traits and ideally including other biodiversity facets that may be affected by insularity, such as phylogeny. Ecological and historical settings across habitats and regions may also provide differences in the insularity forcing, although our results suggest that some persistence traits respond in a similar way across different study systems.

## Conclusions

The functional island biogeography approach applied here allowed us to detect the effect of insularity on biota as well as being useful in capturing varying degrees of insularity. Indeed, we can infer that the strongest effect of insularity on functional traits was found in fens, implying that this system operates more similarly to true islands than outcrops and mountaintops do. This was exemplified by the number of significant trait-insularity relationships, the strength of these relationships, and the differences with matrix-derived species patterns, which were always stronger and more marked in fens than in outcrops and mountaintops (similarly to what was found for specialist species richness in Mendez-Castro et al., in press). Our findings also contribute to the ongoing debate on whether terrestrial island-like systems (including edaphic islands) should be considered insular like true islands, namely whether they are ruled by comparable eco-evolutionary forces as true islands (Itescu, 2019; Jiménez-Alfaro et al., 2021; Patiño et al., 2017). Our work consistently pinpoints how insularity may affect plant species diversity, distribution, and forms not only on true islands (Burns, 2019; Carlquist, 1974) but also in terrestrial-island like systems (such as the studied edaphic islands), and can determine plant abilities to persist locally.

Ultimately, we highlight that insularity – defined as a single (island size, isolation) or combined predictor (target effect) – may operate like a forcing process selecting for adaptive strategies (in terms of trait values and diversity) tailored to promote species ability to avoid local extinction. This biogeographic process may therefore have important implications for the eco-evolutionary dynamics and for the understanding of insular systems’ functioning, species coexistence and evolution. We encourage further studies to implement similar functional island biogeographic approaches, possibly extending to other systems (e.g. true oceanic islands) and focusing on different traits (e.g. related to resource acquisition and dispersal ability; Negoita et al., 2016; Schrader et al., 2021; Taylor et al., 2019).

## Authors’ contribution

GO and LC conceived the research idea and led the writing; MC, MHá, MHo, BJA and DZ provided the vegetation and floristic data and categorised species (specialists and generalists); FEMC measured the insularity metrics; LC gathered plant functional trait data and conducted all statistical analyses; all authors commented on the methodological approach and the manuscript.

## Acknowledgements

LC, FEMC and GO were supported by the Czech Science Foundation (GAČR Project 19-14394Y). MHo was supported by the Czech Science Foundation project no. 19-01775S. MC and MHá were supported by the Czech Science Foundation project 19-28491X. LEMC, LG, JC and GO were also supported by the long-term research development project No. RVO 67985939 of the Czech Academy of Sciences.

## Data accessibility statement

Data (insularity metrics and functional indices) will be made fully available through a public repository such as dryad after manuscript acceptance.

