## Supporting Information for "Insularity forcing on plant persistence strategies in edaphic island systems"

Supporting Information to the paper Conti *et al.* Insularity forcing on plant persistence strategies in edaphic island systems

**Appendix S1:** Species data summary and trait completeness by study system

We provide a table with the trait coverage in each system (i.e. the percentage of species for which the trait information is available; Table S1.1). We also provide a species list for each system, indicating the specialists and the clonal species, in a separated file (Table S1.2).

**Table S1.1.** Number of specialists and total number of species for each dataset, as well as proportion of species in each dataset for which trait data is available

|  | **Fens** | | **Outcrops** | | **Mountaintops** | |
| --- | --- | --- | --- | --- | --- | --- |
|  | **Specialists** | **Total** | **Specialists** | **Total** | **Specialists** | **Total** |
| Number of species | 57 | 317 | 29 | 204 | 42 | 167 |
| Trait |  |  |  |  |  |  |
| Raunkiaer life form | 0.93 | 0.93 | 0.93 | 0.98 | 1.00 | 1.00 |
| Reproduction type | 0.86 | 0.91 | 0.83 | 0.94 | 1.00 | 1.00 |
| Seed mass | 0.74 | 0.81 | 0.76 | 0.91 | - | - |
| Size of the belowground bud bank | 0.81 | 0.88 | 0.83 | 0.92 | - | - |
| Depth of the belowground bud bank | 0.81 | 0.86 | 0.83 | 0.88 | - | - |
| Type of clonal growth organ | 0.74 | 0.71 | 0.38 | 0.40 | - | - |
| Persistence of the clonal growth organ | 0.74 | 0.71 | 0.38 | 0.39 | - | - |
| Lateral spread | 0.74 | 0.71 | 0.38 | 0.40 | - | - |

**Appendix S2:** Models related to matrix-derived species

The same models that were fitted for the set of functional metrics related to the specialist species were fitted also for the metrics based on matrix-derived species. Here we present the results of these models. Generally, the significance and variability explained by the relationships were lower in these models than in models for specialists.

**Table S2.1.** Model outputs for proportion of clonal matrix-derived species in the edaphic island systems. Each column represents a different model with a single insularity metric as predictor. Bold text indicates a relevant model (significant coefficient and/or R^2^ ≥ 0.10). All predictors are mean-centered and scaled by 1 standard deviation. *** p < 0.001; ** p < 0.01; * p < 0.05.

|  | **FENS** | | | **OUTCROPS** | | | **MOUNTAINTOPS** | | |
| --- | --- | --- | --- | --- | --- | --- | --- | --- | --- |
|  | **Model Island size** | **Model Isolation** | **Model Target effect** | **Model Island size** | **Model Isolation** | **Model Target effect** | **Model Island size** | **Model Isolation** | **Model Target effect** |
| Coefficent | **5.61 ***** | **-3.33*** | **-4.84***** | **6.57 ***** | **-4.48*** | **-6.53 ***** | 2.78 | **4.08**** | -0.81 |
| Standard Error | (1.21) | (1.37) | (1.27) | (1.36) | (1.78) | (1.37) | (1.56) | (1.43) | (1.65) |
| R^2^ | **0.32** | **0.11** | **0.24** | **0.56** | **0.26** | **0.56** | **0.12** | **0.26** | **0.01** |

**Table S2.2.** Model outputs for mean trait values at island level considering the matrix-derived species present in the edaphic island systems. Each column represents a different model with a single insularity metric as predictor. Bold text indicates a relevant model (significant coefficient and/or R2 ≥ 0.10). All predictors are mean-centered and scaled by 1 standard deviation. *** p < 0.001; ** p < 0.01; * p < 0.05.

|  | **FENS** | | | **OUTCROPS** | | |
| --- | --- | --- | --- | --- | --- | --- |
| *Seed mass* | **Model Island size** | **Model Isolation** | **Model Target effect** | **Model Island size** | **Model Isolation** | **Model Target effect** |
| Coefficient | 0.01 | -0.05 | -0.04 | **0.37 *** | -0.16 | -0.28 |
| Standard Error | (0.12) | (0.12) | (0.12) | (0.17) | (0.18) | (0.18) |
| R^2^ | 0.00 | 0.00 | 0.00 | **0.21** | 0.04 | **0.12** |
| *Size of the belowground bud bank* |  |  |  |  |  |  |
| Coefficient | 0.15 | -0.02 | -0.08 | 0.20 | -0.07 | -0.14 |
| Standard Error | (0.10) | (0.10) | (0.10) | (0.53) | (0.53) | (0.53) |
| R^2^ | 0.042 | 0.00 | 0.01 | 0.01 | 0.00 | 0.00 |
| *Depth of the belowground bud bank* |  |  |  |  |  |  |
| Coefficient | 0.05 | 0.05 | -0.03 | -0.07 | 0.08 | 0.10 |
| Standard Error | (0.03) | (0.03) | (0.03) | (0.11) | (0.11) | (0.11) |
| R^2^ | 0.04 | 0.01 | 0.02 | 0.02 | 0.03 | 0.04 |
| *Persistence of the clonal growth organ* |  |  |  |  |  |  |
| Coefficient | -0.02 | -0.01 | 0.00 | -0.04 | -0.03 | -0.01 |
| Standard Error | (0.02) | (0.02) | (0.02) | (0.04) | (0.04) | (0.04) |
| R^2^ | 0.01 | 0.00 | 0.00 | 0.07 | 0.04 | 0.01 |
| *Lateral spread* |  |  |  |  |  |  |
| Coefficient | -0.00 | 0.00 | 0.00 | 0.07 | 0.01 | -0.02 |
| Standard Error | (0.02) | (0.02) | (0.02) | (0.04) | (0.04) | (0.04) |
| R^2^ | 0.00 | 0.00 | 0.00 | **0.14** | 0.00 | 0.01 |

**Table S2.3.** Model outputs for diversity of trait values at island level considering the matrix-derived species present in the edaphic island systems. Each column represents a different model with a single insularity metric as predictor. Bold text indicates a relevant model (significant coefficient and/or R^2^ ≥ 0.10). All predictors are mean-centered and scaled by 1 standard deviation. *** p < 0.001; ** p < 0.01; * p < 0.05.

|  | **FENS** | | | **OUTCROPS** | | | **MOUNTAINTOPS** | | |
| --- | --- | --- | --- | --- | --- | --- | --- | --- | --- |
| *Raunkiaer life form* | **Model Island size** | **Model Isolation** | **Model Target effect** | **Model Island size** | **Model Isolation** | **Model Target effect** | **Model Island size** | **Model Isolation** | **Model Target effect** |
| Coefficient | -0.00 | 0.00 | 0.00 | -0.00 | -0.00 | 0.00 | -0.00 | **0.03 ***** | **-0.02 *** |
| Standard Error | (0.01) | (0.01) | (0.01) | (0.01) | (0.01) | (0.01) | (0.01) | (0.01) | (0.01) |
| R^2^ | 0.00 | 0.00 | 0.00 | 0.01 | 0.00 | 0.00 | 0.00 | **0.42** | **0.20** |
| *Seed mass* |  |  |  |  |  |  |  |  |  |
| Coefficient | -0.01 | -0.10 | -0.07 | 0.32 | -0.16 | -0.27 |  |  |  |
| Standard Error | (0.33) | (0.33) | (0.33) | (0.37) | (0.38) | (0.38) |  |  |  |
| R^2^ | 0.00 | 0.00 | 0.00 | 0.04 | 0.01 | 0.03 |  |  |  |
| *Size of the belowground bud bank* |  |  |  |  |  |  |  |  |  |
| Coefficient | 0.49 | -0.33 | -0.45 | -0.36 | 0.33 | 0.44 |  |  |  |
| Standard Error | (0.27) | (0.28) | (0.27) | (0.27) | (0.27) | (0.26) |  |  |  |
| R^2^ | 0.07 | 0.03 | 0.06 | 0.10 | 0.08 | 0.13 |  |  |  |
| *Depth of the belowground bud bank* |  |  |  |  |  |  |  |  |  |
| Coefficient | 0.07 | -0.06 | -0.07 | -0.03 | 0.10 | 0.10 |  |  |  |
| Standard Error | (0.07) | (0.07) | (0.07) | (0.06) | (0.05) | (0.05) |  |  |  |
| R^2^ | 0.017 | 0.014 | 0.02 | 0.019 | **0.18** | **0.19** |  |  |  |
| *Type of clonal growth organ* |  |  |  |  |  |  |  |  |  |
| Coefficient | **0.02 ***** | **-0.01 *** | **-0.02 **** | 0.00 | -0.00 | -0.00 |  |  |  |
| Standard Error | (0.01) | (0.01) | (0.01) | (0.00) | (0.00) | (0.00) |  |  |  |
| R^2^ | **0.21** | **0.11** | **0.20** | 0.05 | 0.02 | 0.05 |  |  |  |
| *Persistence of the clonal growth organ* |  |  |  |  |  |  |  |  |  |
| Coefficient | 0.03 | 0.01 | -0.01 | 0.05 | -0.04 | -0.05 |  |  |  |
| Standard Error | (0.03) | (0.03) | (0.03) | (0.05) | (0.05) | (0.05) |  |  |  |
| R^2^ | 0.03 | 0.00 | 0.00 | 0.07 | 0.03 | 0.06 |  |  |  |
| *Lateral spread* |  |  |  |  |  |  |  |  |  |
| Coefficient | 0.02 | -0.01 | -0.02 | **0.07 *** | -0.05 | **-0.07 *** |  |  |  |
| Standard Error | (0.02) | (0.02) | (0.02) | (0.03) | (0.03) | (0.03) |  |  |  |
| R^2^ | 0.03 | 0.01 | 0.02 | **0.29** | **0.13** | **0.28** |  |  |  |
| *Multiple trait* |  |  |  |  |  |  |  |  |  |
| Coefficient | 0.01 | -0.00 | -0.00 | **0.01 *** | 0.00 | -0.00 |  |  |  |
| Standard Error | (0.00) | (0.00) | (0.00) | (0.00) | (0.00) | (0.00) |  |  |  |
| R^2^ | 0.06 | 0.00 | 0.02 | **0.23** | 0.00 | 0.02 |  |  |  |

**Appendix S3:** Correlation tests on insularity metrics for each system

**Table S3.1.** Spearman rho correlations between insularity metrics. Significant correlations are shown with *** p < 0.001; ** p < 0.01; * p < 0.05.

| **FENS** | Island size | Isolation | Target effect |
| --- | --- | --- | --- |
| Island size | - | -0.49** | -0.84*** |
| Isolation |  | - | 0.86*** |
| Target effect |  |  | - |
| **OUTCROPS** | Island size | Isolation | Target effect |
| Island size | - | -0.01 | -0.49* |
| Isolation |  | - | 0.85** |
| Target effect |  |  | - |
| **MOUNTAINTOPS** | Island size | Isolation | Target effect |
| Island size | - | -0.32 | -0.84*** |
| Isolation |  | - | 0.75** |
| Target effect |  |  | - |
